# Tractable models of ecological assembly

**DOI:** 10.1101/2020.09.02.279943

**Authors:** Carlos A. Serván, Stefano Allesina

## Abstract

Ecological assembly, the way natural communities form under ecological time-scales, is a fundamental and yet poorly understood process. Recent theoretical and empirical approaches to assembly consider systems in which a group of species is introduced in a new environment, and dynamics prune the system down to a sub-community of coexisting species. This “top-down” assembly approach contrasts with the well-studied “bottom-up”, or sequential, assembly, in which species from a pool enter the system one at a time, giving rise to priority effects and complex dynamics. Here we determine under which conditions the two approaches are equivalent, i.e., lead asymptotically to the same exact set of coexisting species. To achieve this result, we represent the assembly process as a network in which nodes are sub-communities and edges stand for invasions shifting the composition of the ecological community from a stable configuration to another. This abstraction makes it easy to determine which states the community can occupy, as well as highlight the potential for priority effects or cyclic species composition. We discuss how the equivalence between bottom-up and top-down assembly can advance our understanding of this challenging process from an empirical and theoretical point of view, informing the study of ecological restoration and the design and control of ecological communities.

## Introduction

Understanding how biodiversity emerges and is maintained is a key challenge in community ecology. When considering a local community and ecological time-scales, this process is called “ecological assembly”, and is realized through the interplay between invasions of species from outside the system, and the interactions between the resident species, driving the community towards a state in which a certain subset of species coexist. In the early days of ecology, assembly—then called succession (Cowles, 1899)—was envisioned as an orderly and predictable process by which the community progressed towards a climax, thereby maximizing some ecosystem function (Cowles, 1899; Odum, 2014).

In the intervening decades, the study of assembly has unveiled a much more complicated picture (Drake, 1991; Law & Morton, 1993; Schreiber & Rittenhouse, 2004; Warren *et al*., 2003)—in general, assembly is anything but predictable: small changes to the order, size, and timing of invasions can result in completely different local communities (Drake, 1991). The dependency of the final community on the order of arrival of invaders gives rise to “historical contingencies” driven by priority effects (Fukami, 2015), and much effort went into determining whether and when these alternative histories can emerge (Fukami & Nakajima, 2011; Warren *et al*., 2003; Zhao *et al*., 2020). Similarly, the density at which the invader enters the community can influence the outcome, a complication compounded by the fact that the local community could be coexisting at a limit-cycle or chaotic attractor—and as such the invader could establish at certain times, but not others.

### Top-down and bottom-up assembly

The advent of cheap sequencing techniques brought the empirical study of the assembly of microbial communities within reach. In this area, a popular experiment is to take the microbial community from an environmental sample and challenge it with a novel, synthetic environment (e.g., grow the community in batch laboratory culture with serial dilution), following the dynamics by which the original community loses members, eventually settling into a sub-community of coexisting species (Bittleston *et al*., 2019; Goldford *et al*., 2018). Interestingly, recent theoretical advances considered the same exact scenario (Barbier *et al*., 2018; Biroli *et al*., 2018; Bunin, 2017; Serván *et al*., 2018) because of its analytical tractability. In abstract, a set of *n* species is introduced (at the same time) in the new environment, and assembly consists of the dynamical pruning of the community down to a set of coexisting species—we therefore term this process “top-down” assembly.

While top-down assembly is realistic for microbial communities (e.g., newborns being colonized by the microbial community of the birth canal), the process seems quite distant from what we expect for macroscopic species. As Karl Sigmund put it “Mother Nature, of course, does not assemble her networks by throwing *n* species in one go. It makes more sense to assume that she adds one species after another through successive invasions” (Sigmund, 1995). Traditionally, this sequential, or “bottom-up” assembly has been the focus of both theory (Capitán *et al*., 2009; Hang-Kwang & Pimm, 1993; Law & Morton, 1993, 1996) and experiments (Drake, 1991; Warren *et al*., 2003).

Clearly, assembling an ecological community through a top-down or bottom-up process could lead to completely different results, even when considering the same pool of species. Because however top-down assembly offers obvious logistical and theoretical advantages, here we ask under which conditions (i.e., models, parametrizations) the two processes are equivalent—they eventually lead to exactly the same final community of coexisting species. Under these conditions, a single experiment could shed light on all possible assembly histories, and analytical results would be within reach.

We first discuss what makes the theoretical study of assembly challenging, and then make a series of assumptions to side-step these difficulties—constraining the problem to make it tractable and yet not trivial. Under these assumptions, we show that the issue of determining whether bottom-up and top-down assembly are equivalent can be turned into a network problem, by representing all possible assembly histories through an “assembly graph” (Capitán *et al*., 2009; Hang-Kwang & Pimm, 1993; Law & Morton, 1993; Schreiber & Rittenhouse, 2004) in which the possible sub-communities we can form from the pool are nodes, and edges stand for invasions that alter the state of the community.

## Methods

We consider a fixed pool of *n* species, and the state of the community through assembly as the set of species present at a given time (the 2^*n*^ sets ranging from ∅, the bare environment, to *{x*_1_, *x*_2_, *…, x*_*n*_*}*, the full species pool), along with their densities.

We model the assembly process of a local ecological community by considering the sequential arrival of invaders from the fixed regional species pool. While easily stated, the study of this problem in full generality is complicated by three main factors.

### What makes the study of assembly challenging?

#### Timing of the invasion events

The time at which the first species goes extinct after an invasion influences the effect of subsequent invasions. For example, take a “rock-paper-scissors” community with three species *{x*_1_, *x*_2_, *x*_3_ *}* (Allesina & Levine, 2011). In this case, the three species can coexist, but no pair of species can. We have that if we start with *{x*_1_, *x*_2_*}*, the resulting community is *{x*_2_*}*; similarly, *{x*_2_, *x*_3_*} → {x*_3_*}*, and *{x*_1_, *x*_3_*} → {x*_1_*}*. Consider the case in which we start with a bare environment (∅), and we introduce species *x*_1_, which can grow in isolation. As such, we have ∅ *→ {x*_1_*}*. Now we introduce species *x*_2_, which would eventually drive *x*_1_ to extinction *{x*_1_, *x*_2_*} → {x*_2_*}*. If we introduce *x*_3_ after *x*_1_ has gone extinct we will have *{x*_2_, *x*_3_*} → {x*_3_*}*; however, if *x*_3_ invades before this happens, then we recover the full community *{x*_1_, *x*_2_ *} → {x*_1_, *x*_2_, *x*_3_*}*. As this simple example highlights, if the speed at which the dynamics of the local community proceed are slow enough compared to the rate of invasion, we have that several species can invade before the community has reached its asymptotic configuration. At the extreme where local dynamics are fast compared to the rate of invasion, we have that each invader finds the local community at its asymptotic state; as the invasion rate increases, the system approaches a point where all the species enter the system before any extinction takes place. If an attractor is reached, then the system conforms to the top-down assembly regime. Increasing the invasion rate even further would result in an open system with constant immigration.

#### Density of the invader at the time of invasion

Consider the two-species competitive Lotka-Volterra model with preemptive competition, and suppose that initially we have species *x*_1_ resting at its carrying capacity, i.e., the state of the system is *{x*_1_*}*. If *x*_2_ invades with sufficiently low density, we find *{x*_1_, *x*_2_*} → {x*_1_*}*; on the other hand, if *x*_2_ has sufficiently high density, we can cross the separatrix in the phase plane, leading to *{x*_1_, *x*_2_*} → {x*_2_*}*. This simple example shows that the density at which the invader enters the system can alter the outcome of the dynamics.

#### Type of attractor

When the local community coexists at a non-fixed point attractor, the fate of the invader could be very different depending on when it is introduced. For example, a predator requiring its prey to be above a certain level would not be able to invade an oscillating system whenever prey are at low abundance, but would start growing if the invasion happened at a time when prey were abundant.

### Ecological assembly without tears

The three features above make the study of assembly in full generality very challenging. To make the problem tractable, here we concentrate on an assembly process that sidesteps these difficulties, and yet is complex enough to generate relevant results.

#### Invasion events are rare

We assume that the invasion rate is low enough such that, after an invasion, the local community has sufficient time to reach its asymptotic configuration before the next invader arrives. In other words, we consider cases in which local dynamics operates at a much faster rate than invasions. Note that this choice precludes certain dynamics; for example, under these stringent conditions the rock-paper-scissors community described above would never reach the three-species configuration. While this is a strong requirement, it corresponds to assumptions routinely made in the study of population genetics (where often only the wildtype and a single mutant interact, Charlesworth & Charlesworth, 2010), adaptive dynamics (Diekmann, 2002), and invasion analysis (Adler *et al*., 2007; Chesson, 2008).

#### Invaders arrive at low abundance

We assume that the density of the invader is low enough so that intraspecific effects are negligible at the time of invasion (again, as routinely assumed in invasion analysis). Under this assumption, the assembly of the Lotka-Volterra preemptive competition model would have two possible final states, corresponding to each species in isolation. Note also that whenever the invader can enter the system only at low abundance, the local stability of an attractor (i.e. the community at the attractor is resistant to small perturbations caused by changes in abundance of any of the species in the pool) is sufficient to make it a possible outcome of the assembly process.

#### Fixed-point attractors

Finally, we consider models in which the asymptotic state of the local community is a feasible, stable equilibrium (we relax this assumption in the Appendix), thereby sidestepping the difficulty stemming from the timing of invasion.

To complement our analysis, we also consider top-down assembly, which violates the first two assumptions above. In this scenario, invasion rate is high-enough that all the species in the pool can attempt to invade before any extinction takes place. As such, the process is equivalent to the scenario in which we initialize the local community with all the species present, at an arbitrary initial condition. Because we retain the third condition, the final community (“assembly endpoint”) represents the attractor reached after the local dynamics have elapsed. Note that also in this case the local stability of an equilibrium is sufficient to make it a possible endpoint of the assembly process.

### Assembly Graph

When the assumptions above are satisfied, we can represent the assembly process as a graph, in which the nodes (vertices) are states of the community, and edges (links) are invasion events, shifting the community from one state to another (see Fig. 1). This assembly graph (Capitán *et al*., 2009; Hang-Kwang & Pimm, 1993; Law & Morton, 1993; Schreiber & Rittenhouse, 2004) recapitulates all possible assembly histories for a pool of species, and its properties determine the outcomes of assembly.

**Fig. 1.**
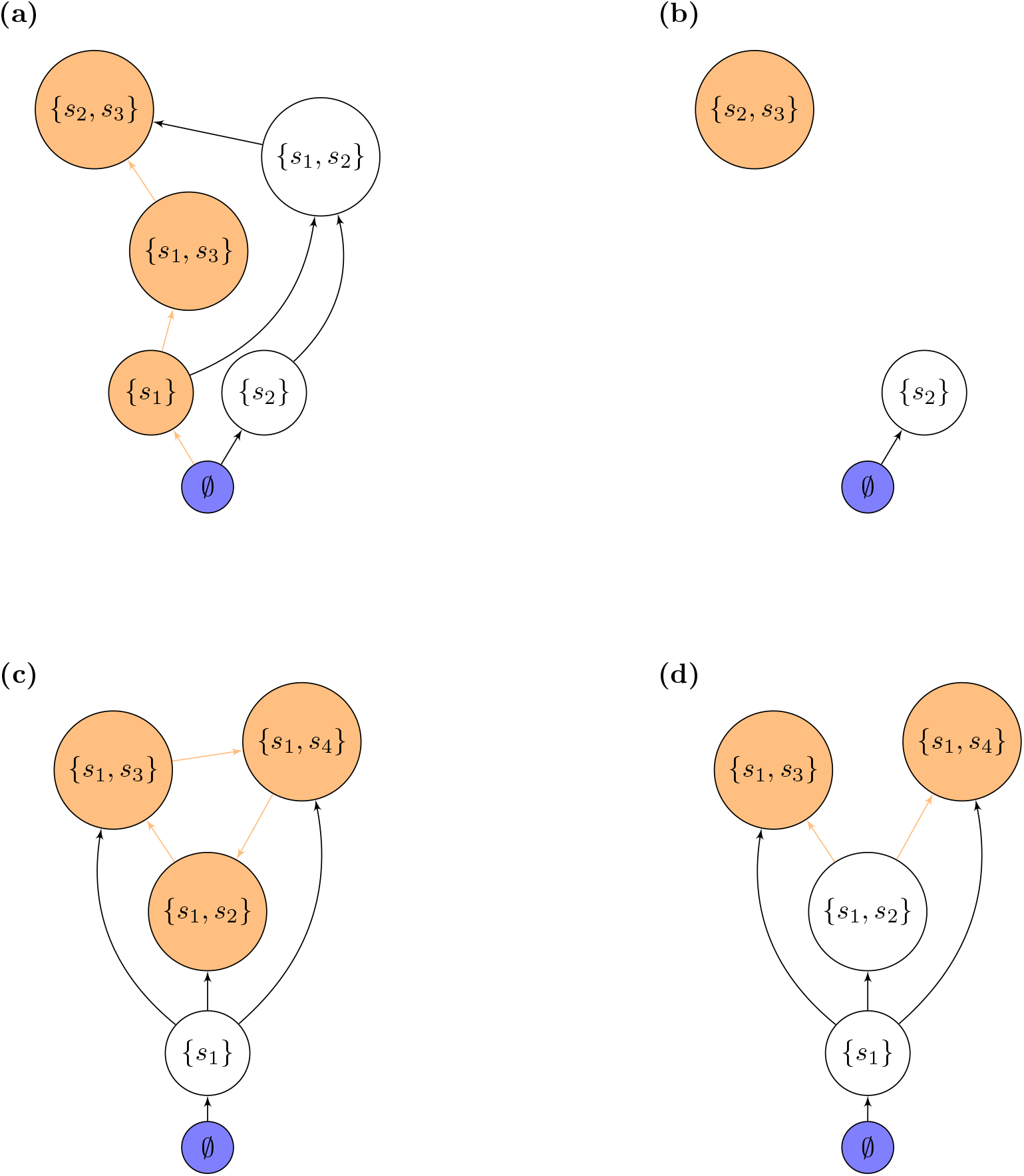
Equivalence between properties of assembly and properties of the assembly graph. Each of the panels shows an hypothetical assembly graphs for a pool of up to four species. **(a)** This assembly graph is accessible—i.e., any of the states can be reached through a walk starting at ∅. A possible assembly path leading to *{s*_2_, *s*_3_*}* is highlighted in orange. Note that *{s*_2_, *s*_3_*}* is the unique sink for the graph, meaning that asymptotically assembly will always result in this community. In order to reach the final state, species *s*_1_ needs to be present at some point of the assembly history. If the species *s*_1_ is not available in the pool, we obtain **(b)**, which is not accessible: the state *{s*_2_, *s*_3_*}* can be reached by top-down, but not bottom-up assembly, while bottom-up assembly will always result in *{s*_2_*}*. This graph possesses two separate endpoints. The graph **(c)** contains a more complex sink (highlighted in orange) composed of a sequence of three communities connected by a cycle. Assembly proceeds from the empty state to *{s*_1_*}*, and from here is trapped in the cycle. Finally, **(d)** represents a graph that is accessible, but has two possible endpoints, giving rise to “historical contingencies”.

*G* is a directed graph with nodes *V* (*G*) and edges *E*(*G*). The nodes represent the feasible, stable sub-communities that can be formed from the pool, composed by a certain number of species along with their densities *S* = *{x*_1_, *… x*_*n*_*}* = *x*^*S*^ (where some of the *x*_*i*_ could be zero or, equivalently, absent). If for each set of species we have at most one such state (i.e., the system does not display true multistability), then there are up to 2^*n*^ nodes, and communities are uniquely identified by their composition; if multistability is possible, on the other hand, we can have multiple nodes representing communities with the same set of species, but different densities. The bare environment *{*0, *…*, 0*}* = ∅ is always included in the graph. The edges are determined by invasion events. Let *j* be a species: *j* connects two distinct vertices *S* and *S*^*′*^ in *G* if: i) *j* is not in *S*, ii) *j* can invade *x*^*S*^ when rare (*x*_*j*_ ≈ 0), and, iii) when *j* invades, dynamics push the system to *S*^*′*^. We denote this transition by 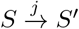.

The explicit separation of timescales between invasions and local dynamics allows the assembly graph *G* to completely capture the assembly process: any sequence of successful invasions will be represented as a walk (i.e., a series of nodes connected by edges) within *G*, and all the possible communities that can be observed are nodes of *G*. Importantly, this means that we can study the assembly process directly on *G*, and forget about the dynamics altogether.

For any set *S* of coexisting species, we can ask whether *S* can arise through bottom-up assembly. In this scenario, we can only observe a subset *S* if there is a sequence of invasions taking the system from the empty state to *S*. Thus, we are interested in determining whether, for any vertex *S*, there is a path in *G* starting at ∅ and ending at *S*. Such a path is called an assembly path for *S*. We call an assembly graph *G* “accessible” if all subsets *S* in *G* have at least one assembly path. If some nodes are not accessible, they can be reached from top-down, but not bottom-up assembly.

Historical contingencies arise if different sequences of invaders can drive the community towards different final states. We call the final states of the assembly process “assembly endpoints” (also known as “permanent” or “persistent” states, Drake, 1991; Hang-Kwang & Pimm, 1993; Law & Morton, 1993; Schreiber & Rittenhouse, 2004; Warren *et al*., 2003). As such, the existence of historical contingencies requires the existence of multiple assembly endpoints.

Here we distinguish between two types of endpoints. In the simplest case, there is a set of species *S* which is resistant to invasions by any other (combination of) species in the pool. Thus *S* is a node with no outgoing edges (called a “sink” in graph theory). Alternatively, an endpoint is comprised of a set of feasible communities *𝒰* = *{S*_1_, *…, S*_*k*_*}* for which invasions only transition the system within the set *𝒰*, and any two sub-communities *S*_*i*_, *S*_*j*_ in *𝒰* are connected by a path in *G*. In this case, rather than having a single-vertex endpoint, we have a set of communities that function as a sink: once the assembly process reaches any of these nodes, the system can only move between the states in *𝒰* (see Appendix S1 for a more detailed discussion).

The more complex endpoint described above always contains an assembly cycle: a set of communities *{S*_1_, *…, S*_*l*_*}* such that we have a sequence of invasions taking *S*_1_ *→… →S*_*l*_ *→S*_1_ (for an example, see Schreiber & Rittenhouse, 2004). Trivially, assembly cycles map to directed cycles (in the graph-theoretical sense) within the assembly graph *G*. For a cycle to exist, it is necessary that at least one invasion event causes an extinction. Suppose that transitions between 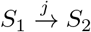 causing extinctions are only possible if *S*_1_ *∪ j* cannot coexist (this will happen for example if feasibility of all the species in a community implies global stability of the attractor). If we further assume that a successful invader is always present in the next state, then cycles are the natural generalization of intransitive competition (e.g., rock-paper-scissors communities). Indeed, for a cycle to occur under these conditions, we need a species *j* and a state *S*_1_ so that *j* drives some of the species in *S*_1_ extinct, and conversely to go back to *S*_1_ there must be a species in the total set of invaders that makes *j* go extinct.

All the properties of the assembly graph are summarized in Fig. 1, showing a collection of hypothetical assembly graphs with different characteristics.

## Results

The assembly graph described above is quite abstract. To provide a concrete example, we show how the graph can be built and analyzed for the Generalized Lotka-Volterra (GLV) model (see the Appendix S5 for a consumer-resource model):

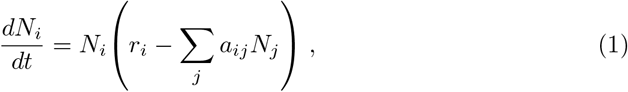

Here, the pool of species is defined by the parameters: the vector of growth rates *r* and the matrix of species interactions *A* = (*a*_*ij*_). For simplicity, we consider the case of competitive communities, in which each species is assigned a positive growth rate *r*_*i*_ *>* 0, so that it can grow in isolation. Next, we need to ensure that the model can only give rise to fixed-point attractors (equilibria), to comply with the assumptions above. To this end, we take the interaction matrix *A* to be of the form *A* = *D*(*v*)*BD*(*w*), where *B* is a non-singular, symmetric matrix, with nonnegative entries, and *D*(*v*) and *D*(*w*) are diagonal matrices with diagonal entries equal to *v* and *w*, respectively. Further, we assume that *v*_*i*_ *>* 0 and *w*_*i*_ *>* 0 for all *i*, ensuring that species compete with each other. Notice that *D*(*v*)*BD*(*w*) can always be expressed such that *b*_*ii*_ = 1, by reabsorbing the diagonal of *B* into *v* and *w*. As such, we only consider matrices with *b*_*ii*_ = 1 for all *i*. The parameter *v*_*i*_ modulates the effect of other species on the growth of species *i*, and is thus related to the resource requirements of *i*. Conversely, *w*_*i*_ models the resource use of species *i*. Eq. (1) becomes:

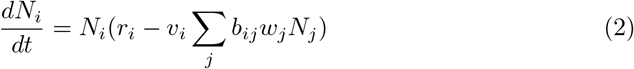

By a change of variables (*w*_*j*_*N*_*j*_ *→ x*_*j*_, *γ*_*i*_ = *r*_*i*_*/v*_*i*_), we obtain:

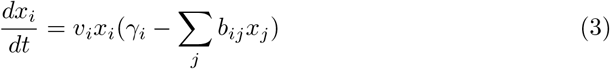

This classic system of equations has been studied for decades, and MacArthur (1970) constructed a global Lyapunov function for the system, which is maximized through the dynamics (see also Appendix S4):

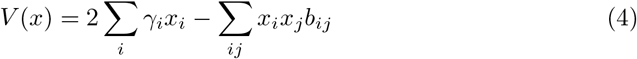

For any equilibrium *x*^*\**^, we find: 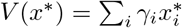. This quantity has a more intuitive interpretation in the case *r*_*i*_ = *v*_*i*_ (and thus *γ*_*i*_ = 1), in which case *V* at equilibrium is simply the total biomass of the system.

Because *B* is non-singular and symmetric, the model yields only fixed-point attractors. If we further take *B* to be stable (i.e., with only positive eigenvalues, and therefore positive definite), we reduce the number of attractors to one (Hofbauer & Sigmund, 1998). In fact, when *B* is stable, the unique attractor for the top-down assembly is characterized by the following: we can identify a set of species *S ⊆ {*1, *…, n}* which can coexist at a positive equilibrium (feasibility) and moreover any species not present in *S* cannot invade when rare (non-invasibility). This type of attractor is called a saturated rest point, or non-invasible solution (Hofbauer & Sigmund, 1998; Serván *et al*., 2018).

Because of the restrictions on Eq. (2), any subset of species will follow the same type of dynamics (i.e., any sub-set of species defines a new pool with the same properties—mathematically, any minor of a positive definite matrix is also positive definite), and thus, if we initialize the system with any subset of species at arbitrary positive initial conditions, dynamics will converge to a unique globally-attractive fixed-point with the same characterization.

This result yields a simple test to assess if the arrival of a new species can push the community to a new state: it is sufficient to determine whether the species can invade when rare. If so, then the resident community, or any of its subcommunities, cannot be the attractor for the augmented community. Thus, the new invader must be part of the new configuration, thereby changing community composition. Consequently, for any invader *j*, we call the invasion successfull if *j* can invade when rare. While rarely formalized, these conditions are implicitly assumed when performing invasion analysis (Adler *et al*., 2007).

When *B* is not stable, the community can have more than one attractor (Biroli *et al*., 2018). Nevertheless, as shown in the Appendix (S4), each attractor *S* is characterized by the same conditions as for the case of *B* stable: the equilibrium with the species *S* is a feasible and non-invasible set of species, whose interaction matrix *B*^(*S*)^ (i.e., the matrix *B* in which we retain only the rows and columns corresponding to the species in *S*) is stable.

### Equivalence between top-down and bottom-up assembly

Having defined the model, we can determine whether top-down (all in one bout invasion) and bottom-up (sequential) assembly are equivalent: *given a species pool, the bottom-up assembly endpoints are the same as the endpoints for the top-down assembly process*. If this is the case, given sufficient time, the bottom-up assembly process will result in one of the endpoints that are accessible via top-down assembly. Because in this case (and whenever this holds) the attractors for top-down assembly are in correspondence with the sink nodes in the assembly graph, the equivalence between assembly process maps to the following graph property: *Every walk in G eventually reaches a sink node*.

First, we consider the case in which *B* is stable (positive definite), for which case we prove (Appendix S2) three results. First, the graph *G* is accessible, i.e., there is a path leading from ∅ to any feasible community *S*. This means that we can potentially build from the bottom-up any subset of the species of the pool which can coexist together. Moreover, for each *S* in *G* we can find an assembly path in which no species goes extinct. As such, to build *S* from the ∅, we only need to choose the right order of invasions within *S*, guaranteeing that we will not observe “Humpty-Dumpty” communities (Law & Morton, 1996; Pimm, 1991; Warren *et al*., 2003). Second, *G* has no (directed) cycles. Third, *G* has unique sink *u* and source ∅. Coupled with the first two properties, this implies that historical contingencies are impossible—asymptotically, the assembly will always result in *u*.

Moreover, for any graph with the above properties, there exists an ordering of the vertices such that two of them are connected only if one appears after the other (i.e., a topological sorting, Cormen *et al*., 2009). This implies that we can find a quantity *Q* that increases monotonically as the assembly process unfolds. Thus, the assembly process “optimizes” *Q*. In the example of Fig. 2, we can take *Q* to be the height of each node in the plot. For the assembly model, *Q* has a concrete meaning: it can be chosen as the Lyapunov function *V* in Eq. (4). In the case of *r*_*i*_ = *v*_*i*_, we have that for each feasible subset *S, V* represents the total biomass at the equilibrium *x*^*S*^. Thus, the assembly endpoint *u* is the feasible sub-community with the maximal total biomass, and at each step along an assembly path the total biomass present in the local community increases (Fig. 2).

**Fig. 2:**
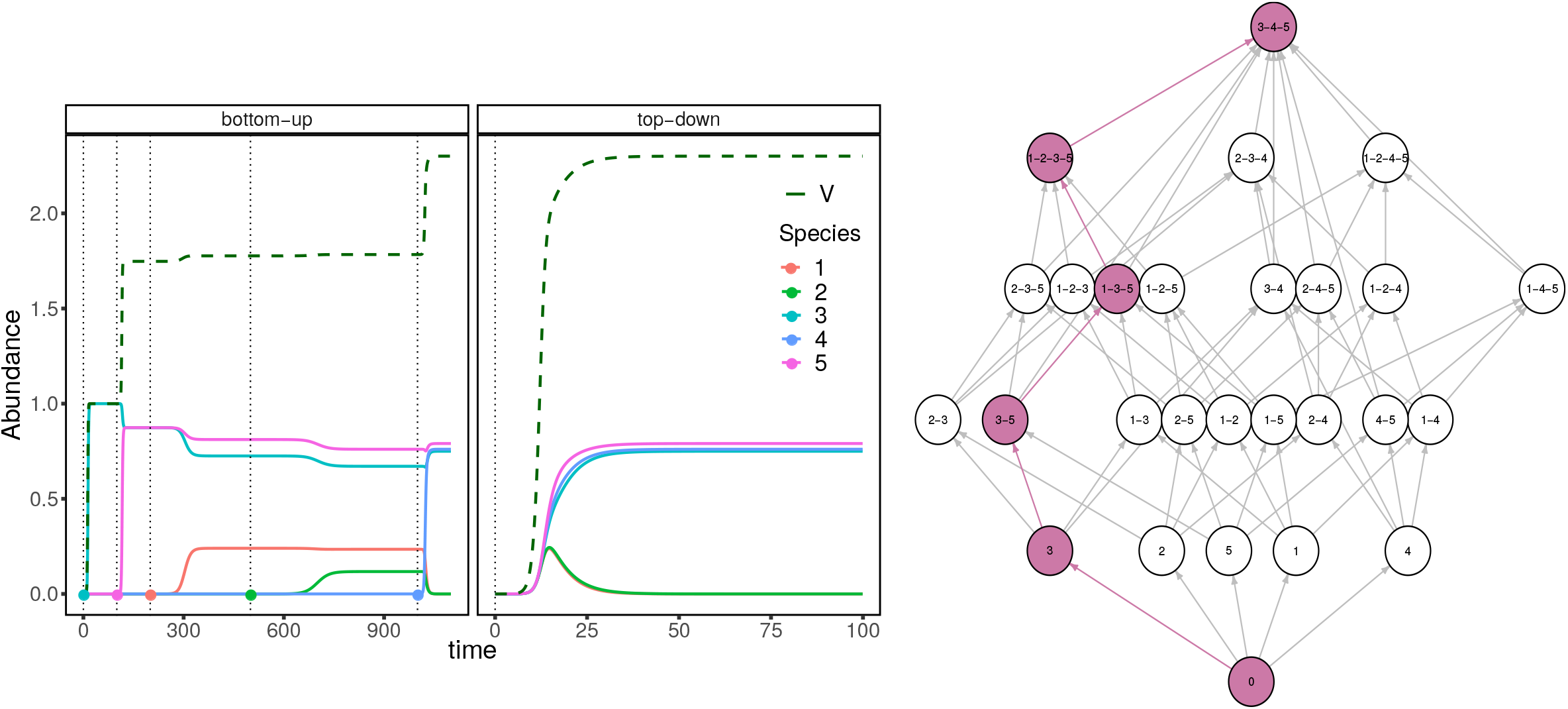
Assembly trajectories (left) and assembly graph (right) for a pool of 5 species, following the GLV model in Eq. 2 with *r* = *v*, and interaction matrix *B* stable. The trajectories are shown for a possible bottom-up assembly sequence (left panel) and for top-down assembly (right panel). Vertical dashed lines show the timing of the introduction of an invader (colored point). For bottom-up assembly, at each invasion the system moves from a stable state to another, as highlighted in the assembly graph on the right. The quantity *V* (dark green dashed line) shows a non-decreasing trajectory for both types of assembly processes, showing that *V* is maximized through assembly (and dynamics).

By the non-invasibility characterization of the attractors in Eq. (2) the unique sink vertex in *G, u*, is precisely the unique attractor of the dynamics for Eq. (2). That is, *u* corresponds to the state that would be reached when we initialize the system will all species present (top-down assembly). Because of the fact that *G* is accessible, *u* can be observed during assembly. Finally, acyclicity of *G* rules out the possibility of other types of assembly endpoints. Thus, it follows that when *B* is stable, the top-down and bottom-up assembly processes are equivalent.

When we relax the stability condition for *B*, there is no guarantee that a unique sink in the graph will exist (Biroli *et al*., 2018). In this setting, therefore, priority effects and historical contingencies might play a role. Yet, a global Lyapunov function *V* for Eq. (2) exists regardless of the stability of the interaction matrix *B* (MacArthur, 1970). *As for the stable case, the existence of V* implies that *G* is acyclic. Similarly, the attractors of the dynamics satisfy the same non-invasibility conditions, akin to the case of *B* stable (see Appendix S4). Thus, regardless of the stability of *B*, the assembly graph *G* is accessible and acyclic, with sink vertices in correspondence to attractors for the top-down assembly process. Hence, bottom-up and top-down assembly are still equivalent: there are initial conditions for the top-down scenario leading to the same exact endpoints as found in the bottom-up scenario with different invasion sequences (Fig. 3).

**Fig. 3:**
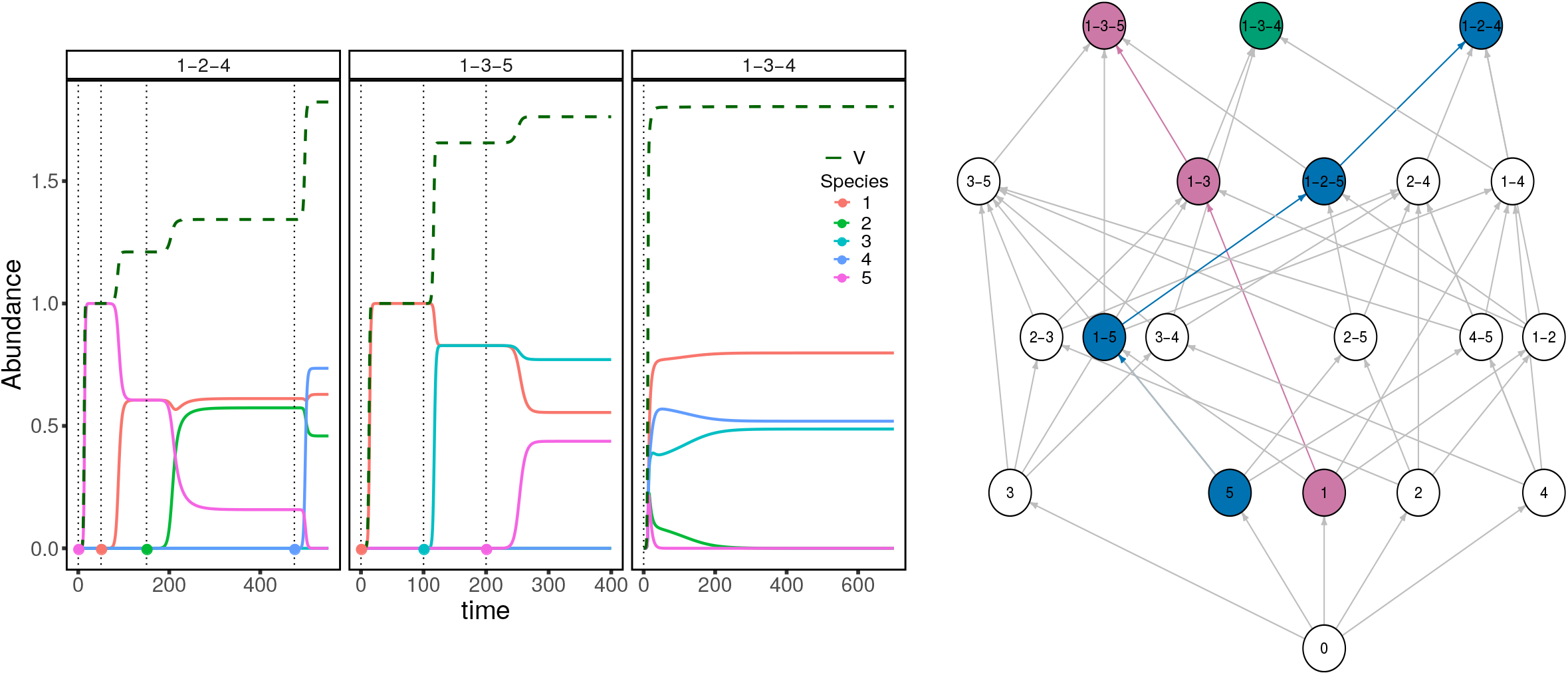
As Fig. 2, but with an unstable matrix of interactions *B*, leading to priority effects and historical contingencies. The trajectories on the left are labeled according to the assembly endpoint they reach. In the middle and left panel we proceed by bottom-up assembly, and the sequence of invasions is highlighted in the assembly graph on the right. In the rightmost panel we start with all the species at low density and converge to the subset *{*1, 3, 4*}*. The quantity *V* is plotted in dashed dark green and again shows a non-decreasing trajectory in all three different assembly processes.

In summary, for the competitive GLV model in Eq. (2), we have that bottom-up and top-down assembly are equivalent: given sufficient time, bottom-up assembly will result in the same community (when *B* is stable) or in one of the several communities (when *B* is unstable) that can be reached by choosing appropriate initial conditions in a top-down assembly. Importantly, in both cases the assembly graph is accessible (i.e., all possible communities and coexisting species can be formed by sequential invasions) and acyclic (i.e., the endpoint of assembly are given by specific communities, rather than ensembles of communities connected by cycles). These properties guarantee that, under these conditions, assembly is indeed an orderly, predictable process, as envisioned by the pioneers of succession. In fact, we can even pinpoint the exact ecosystem function being maximized through dynamics and assembly (e.g., biomass for the simplest choice of parameters).

In the Appendix (S2), we show that, more generally, whenever assembly maximizes a given function, then top-down and bottom-up assembly are equivalent. Less stringently, if we want to recover only the accessibility of *G* (i.e., want to know any feasible subcommunity can be observed during bottom-up assembly), it is sufficient to impose optimization only locally (i.e., a function is maximized when assembling using each feasible subset of the pool). In this case, however, cycles might be present (Appendix S2).

## Discussion and Conclusion

Inspired by recent experiments in microbial ecology (Bittleston *et al*., 2019; Goldford *et al*., 2018), as well as theoretical developments (Barbier *et al*., 2018; Biroli *et al*., 2018; Bunin, 2017; Serván *et al*., 2018), we have attempted connecting the “top-down assembly” process, in which all species from a pool are introduced in an environment at the same time, with the “bottom-up assembly” process, in which species invade the system one at a time. To draw this connection, we sidestepped some of the main obstacles to the development of a theory of ecological assembly, by assuming that invasions are rare, invaders enter the system at low abundance, and that each community settles into a stable equilibrium. Under these conditions, all possible assembly histories one can form using a pool of species can be summarized in an assembly graph (Capitán *et al*., 2009; Hang-Kwang & Pimm, 1993; Law & Morton, 1993; Schreiber & Rittenhouse, 2004), which then can be analyzed directly, foregoing dynamics completely.

We concentrated on a few simple properties of the graph: if a node is reachable starting from the source (∅), then the corresponding community can be assembled bottom-up (accessibility); if the graph is acyclic, then there is a natural ordering of the communities, in line with the pioneering ideas about succession (Cowles, 1899; Odum, 2014); historical contingencies can only play a role whenever there are multiple sinks. In the case in which the graph is accessible, acyclic and has a single sink, then there exists a function that is maximized through assembly, and all assembly histories will eventually lead to the same community.

As an application, we showed that for the symmetric Lotka-Volterra model this framework allows us to prove the equivalence of top-down and bottom-up assembly: the communities that are reached asymptotically through either sequential invasions or a single, massive invasion are exactly the same. The condition for this equivalence is that assembly proceeds as an optimization process. However, this equivalence—as we saw when relaxing the stability assumption for the local dynamics—does not preclude the existence of multiple assembly endpoints. Thus, while assembly may appear orderly and predictable along a particular assembly trajectory, the whole process is in general susceptible to priority effects and historical contingencies. Similar conclusions are reached for a consumer-resource model (Appendix S5).

Our framework can be expanded in a variety of ways. For example, in the Appendix (S3) we relax the assumption that invasions are spaced apart far enough to allow the community to settle into its asymptotic state before the next invasion. In this way, multiple invasions can interact with each other (reminiscent of multiple competing mutations in population genetics, Messer *et al*., 2016).

Connecting top-down and bottom-up assembly can help with the development of ecological applications and experimental approaches across several domains.

For example, it validates the use of top-down assembly (which by definition can lead to any of the feasible sub-communities) in experiments. Suppose that the top-down assembly of a pool of species, performed at different initial conditions, yields markedly different communities—this would suggest that the assembly graph has multiple sinks, and as such the pool of species would be a good candidate for the study of priority effects. If, on the other hand, top-down assembly were to always lead to the same outcome, then the case for alternative assembly histories would be considerably weakened.

Similarly, assembly is key for ecological restoration. Suppose one wants to restore an ecological community, and assume that the desired, pristine community was an assembly endpoint. Even having access to all the members of the desired community might not be sufficient to restore the system, unless the assembly graph contains an assembly sequence comprised purely of members of the desired community (Pimm, 1991). This is not the case whenever to achieve a certain state through assembly, we need to go through “stepping stone” communities in which some invaders enter transiently in the system, and then disappear (as found experimentally, Amor *et al*., 2019; Warren *et al*., 2003). In such cases, to bring the system to the desired state *S*, we would need to be able to introduce the species in *S* as well as any other transient invader in the right order—a much more challenging situation. Studying whether and when we can rebuild a community by introducing only the desired species in more complex models would be therefore be of both theoretical and applied interest (Law & Morton, 1996; Schreiber & Rittenhouse, 2004; Temperton *et al*., 2013; Warren *et al*., 2003).

Finally, assembly informs the construction of ecological communities with desired properties (e.g., polycultures of algae for biofuel production, Newby *et al*., 2016). When confronted with a large pool of candidate species to choose from, it is difficult to test all possible communities, and top-down assembly could yield smaller subsets to tinker with.

Our main theoretical device was the translation of the assembly process to a language centered around the assembly graph *G* (Capitán *et al*., 2009; Hang-Kwang & Pimm, 1993; Law & Morton, 1993). In fact, the main reward for the stringent assumptions we made is that in this case *G* captures completely the assembly process—features of the assembly dynamics are reflected in properties of *G*, which is much easier to study. We expect that further investigations of this graph will shed light on other features of assembly. For example, when multiple assembly endpoints exist, what is the probability that the process culminates at a particular endpoint? Studying random walks on *G*, where the structure of *G* determines the transition graph, may provide insights into this question (Capitán *et al*., 2009).

The equivalence we have established is asymptotic in nature. For a large enough species pool, the number of invasions needed to converge to the final endpoint can be very large; in fact, even when there is a unique assembly endpoint, observing communities after a fixed number of steps along distinct assembly sequences may show markedly different compositions. By computing the distribution of the length of the walks between the empty community and the sink nodes in *G*, one estimates the speed of assembly, and gauges the importance of transient assembly dynamics. Similarly, while the absence of cycles implies the equivalence between the two assembly processes, it is not a necessary condition for the equivalence (see Appendix S2 for an example).

Our results suggest that we can approximate the outcomes of bottom-up assembly by analyzing the outcomes of the much more manageable process of top-down assembly. Moreover, we note that the structure of *G* can be studied empirically. Assuming Lotka-Volterra dynamics, the parameters needed to determine all the nodes in *G* are precisely those that can be inferred from coexistence-type experiments (Maynard *et al*., 2020). Importantly, the subset of experiments needed to perform this inference scales linearly with the number of species, opening the door to empirical approaches to build the assembly graph for large communities. These improvements, in combination with the results for top-down assembly, enable the formulation of testable predictions. And, in guiding the design and interpretation of novel experiments, our results will further elucidate the structure of ecological assembly.

We thank J. Capitán, P. Lechón and M.O. Carlson for feedback on the manuscript. Special thanks to J. Grilli and Z. Miller for comments and discussion at different stages of this work. 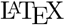 style based on a template by Phillip Schlegel.

### Appendix

Consider a community of *N* = *{*1, *…, n}* species whose dynamics are described by a set of autonomous ODEs:

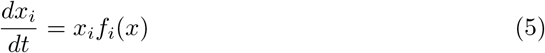

In the following, we explore sufficient conditions under which an assembly process, defined by a pool of species and local dynamics given by Eq. (5), satisfies the equivalence between bottom-up and top-down assembly. To do so, we turn to the study of the properties of the assembly graph *G* described in the main text. Equivalence in this case explicitly means that we can build an assembly sequence starting from the bare environment, and reach each of the configurations that can be observed when starting with all the species present at the same time. In other words, the assembly endpoints are the same for bottom-up assembly and top-down assembly.

### S1 Assembly Graph

#### Definition

For any subset of species *S ⊆ {*1, *…, n}* let Σ_*S*_ be an attractor for the for the dynamics restricted to the species in *S*. We call Σ_*S*_ feasible if Σ_*S*_ contains all the species in *S*. We assume that the following conditions, which hold for the competitive GLV system of the main text, are satisfied by the dynamics given by Eq. (5):

##### Condition 1

a. *Any feasible* Σ_*S*_ *is globally attractive within S. Thus, there is no true multistability (i*.*e*., *any feasible state is uniquely defined by the species it contains);*
b. *Let x*^*S*^ *be the time-average of* Σ_*S*_. *Then, an invasion for j ∉ S is successfull if j can invade the subsystem S at x*^*S*^ *when introduced at low density (x*_*j*_ ≈ 0).

Define the assembly graph *G* in an analogous manner as in the main text:

a. *V* (*G*) is indexed by the feasible sets *S*;
b. An edge connects nodes *S* and *S*^*′*^ if there is a species *j* that can invade *S*, and the dynamics of the augmented system ends up in Σ_*S*_*′*.

For any subset *S*, we denote *G*_*s*_ to be the subgraph of *G* induced by vertices corresponding to sub-communities of *S*.

#### Assembly end-points

An assembly endpoint is the final set of configurations of the local community constructed during assembly (Drake, 1991; Hang-Kwang & Pimm, 1993; Law & Morton, 1993; Schreiber & Rittenhouse, 2004; Warren *et al*., 2003). By definition, we have transitions between states only if a state can be invaded by at least one other species. Thus, assembly endpoints are configurations such that invasions do not take the system out of them. Of course, then, the whole set of communities would be an assembly endpoint! In order to avoid this trivial result, we need to impose a minimality condition: we are only looking for sets that satisfy the previous property and which do *not* include any other such set. We can use the assembly graph *G* to make this statement more precise: a set of nodes *U* = *{S*_1_, *…, S*_*n*_*}* of *G*, is an assembly endpoint if and only if, for any *S*_*i*_, its outgoing links points within *U*, and moreover any such *S*_*i*_ can be reached from another *S*_*j*_ in *U* by a sequence of invasions. Thus *U* is a *strongly connected component* of *G*, and moreover, when collapsing the graph *G* by the equivalence relation of *s ∼ s*^*′*^, if there are directed paths *s → s*^*′*^ and *s*^*′*^ *→ s* in *G, U* is a sink.

### S2 Equivalence

A necessary condition for the equivalence of the two types of assembly is that assembly endpoints are in correspondence with attractors for the dynamics of Eq. (5). Since no assembly endpoint with more than one vertex satisfies such constraint, it follows that, when the two assembly processes are equivalent, all assembly endpoints are comprised of a single vertex. This then forces the attractors to be characterized by a non-invasibility condition:

#### Condition 2

*Let S*^*′*^ *be a sink vertex in G*_*S*_, *then* Σ_*S*_*/ is an attractor for the dynamics of* (5) *restricted to S. Furthermore, any attractor for any subsytem of* (5) *corresponds to such a vertex, i*.*e*., *attractors are characterized by being non-invasible by any other species (one at a time) not present in them*.

In conjuction with condition 1, condition 2 implies that *G*_*S*_ has a unique sink vertex (*S*) for any feasible *S*.

With the correct language now in place, we can state our main result:

##### Theorem 1

a. *Assuming conditions 1 and 2, and G*_*S*_ *acyclic for any feasible subset S, then G is accessible. Furthermore, for any feasible subset S there is an assembly sequence reaching S with no extinctions, and built purely from species present in S*.
b. *If, in addition, G is acyclic, then bottom-up assembly is equivalent to top-down assembly*.

By theorem 1, to determine the equivalence between the assembly scenarios when the local dynamics are described by the GLV model of the main text, it is sufficient to check that the assembly graph *G* is acyclic. As shown in the next section, this is proved by the existence of the Lyapunov function *V*.

#### Acyclicity

As stated in the main text, if a graph *G* is a directed acyclic graph, there exists a quantity *Q* that we can associate to each node such that *Q* increases along assembly paths. In the context of community dynamics, this condition is equivalent to:

##### Condition 3

*For any attractor* Σ_*S*_, *take x*^*S*^ *to be its time average. Then, there is a (smooth) function h* : ℝ^*n*^ *→*ℝ, *such that we can order x*^*S*^ *by h*(*x*^*S*^), *and an edge exist between* Σ_*S*_ *→* Σ_*S*_*′ only if h*(*x*^*S*^) *< h*(*x*^*S*^*′*).

**Lemma 1**. *G is a directed acyclic graph (DAG) if and only if we can find such an h*.

*Proof*. If *G* is a *DAG* (directed acyclic graph), by topological sorting we find a quantity *Q*, and we can use it order the nodes (attractors). Thus *Q* : *V* (*G*) *→* ℕ, and *Q* increases along each edge. The set of equilibria *{x*^*S*^*}* is finite, and thus discrete, and therefore we can cover it by disjoint balls of radius *ε*. Define for each a bump function 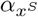 supported at *B*_*ε*_(*x*^*S*^), then define 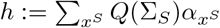.

To prove the converse, suppose to the contrary that there is a loop 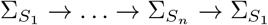. Then by definition of *h* we must have 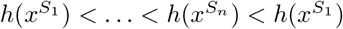—a clear contradiction.

It is important to remark that *h* is *not necessarily a global Lyapunov function* for the dynamics of (5). The existence of a Lyapunov function provides such an *h*, but in general not the other way around. From the biological point of view, the hope would be that *h* can be given a more concrete meaning, as done in the example of the main text where under certain parameterizations *h* is simply the total biomass for the community.

#### Accessibility

Accessibility of *G* implies that any node (besides the empty state) has at least one *incoming* edge, thus the only *source* node of *G* is ∅. If *G* is acyclic, both conditions are equivalent:

**Lemma 2**. *Let G be a* DAG, *then G is accessible if and only if G has a unique source node*.

*Proof*. The preceeding discussion shows that if *G* is accessible, then it has a unique source node.

For the converse, let Σ_*S*_ *≠* ∅ be a node in *G*. Then, by definition we can find an incoming edge 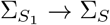, and if 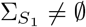, we can repeat the process. If we proceed for long enough, we must eventually find, because otherwise we would have a cycle. Indeed, take a path of length larger than the number of vertices in *G*, then by the pigeonhole principle, two of the nodes must be the same—hence we have a cycle. Since *G* is acyclic this concludes the proof.

We distinguish between two types of incoming links:

a. **additions:** The invasion event does not cause the extinction of any of the species present in the system;
b. **extinctions:** The invasion event causes some of the species in the attractor to go extinct.

If *G* has a unique source node, and at least one of the edges for each node is of the first type, then we can recover a *perfect addition ordering* for each node (Sugihara, 1984). In particular, any attractor can be recovered by assembly *purely from its members*. Furthermore, this implies that for any feasible subset *S*, there is a species *i ∈ S* such that the removal of *i* from *S* does not cause any further extinction.

If, on the contrary, we have a vertex *S* for which all the incoming edges are of the second type, then we must have *stepping-stone* species (Amor *et al*., 2019; Warren *et al*., 2003), meaning that some species are necessary for the assembly process to lead to a given community composition, but are not part of it. And, contrary to the above, the removal of any species from a feasible set *S* will cause an extinction cascade.

The same proof as for lemma 2, but in reverse, shows that:

**Proposition 1**. *If for each feasible subset S, G*_*S*_ *is acyclic and has a unique sink node, then G*_*S*_ *(and therefore G) is accessible. Moreover, we can find a perfect addition ordering for each S*.

The last proposition then implies that we do not need *global* acyclicity for *G* to be accessible, we simply need the graph restricted to feasible sets to be acyclic which is a more *local* condition, especially if the typical behavior of the community is that a considerable fraction of the species cannot coexist (Barbier *et al*., 2018; Biroli *et al*., 2018; Bunin, 2017; Serván *et al*., 2018). It remains to show the last statement of theorem 1.

*Proof of Theorem 1*. The same type of argument as in lemma 2 shows that, if *G* is acyclic, we can always build an assembly sequence such that we reach a sink node—thus, both processes are equivalent since any assembly endpoint with more than one vertex contains a cycle.

### S3 Extensions

Thus far, we have made an explicit separation of timescales between invasion events and local dynamics. This allowed us to only consider the asymptotic behavoir of the local communities along an assembly path. If this condition is not satified, transient dynamics can play an important role (for example, see Schreiber & Rittenhouse, 2004, and recall the rock-paper-scissors example in the main text). To lift this condition, we can model the number of invasion events as a Poisson process with rate *ρ*. At each invasion event, an invader *j* is sampled uniformly at random from the set of species in the pool, and is added to the community at low density, *x*_*j*_ ≈ 0. Let *S*_1_, *S*_2_ be two feasible states, such that asymptotically we have 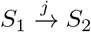. Let *t*_12_ be the time it takes for the community to reach its asymptotic attractor. The probability that such a transition is observed under the new model is simply 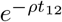. Thus, in the limit of *ρ →* 0 we recover the bottom-up assembly with one invasion at a time (studied in the main text), with an assembly graph denoted by *G*_asym_. In the limit of *ρ→ ∞* all the species are present in the local community, yielding a Lotka-Volterra model with immigration (Biroli *et al*., 2018). To explore how the process changes along *ρ*, we take the assembly graph *G* to be a function of *ρ*.

Let *j*_1_, *j*_2_, *…, j* be a sequence of invaders arriving to the local community at times *t*_1_, *…, t*_*n*_. Let *S*_1_ = *{j*_1_*}*, and *τ*_1_ the time at which the local community reached an asymptotic configuration *S*_2_. This means that all the invaders arriving between *t*_1_ and *τ*_1_ push the system from *S*_1_ *→ S*_2_. Proceeding in this fashion, we have transitions *S*_1_ *→ S*_2_ *→ … → S*_*m*_ at times *τ*_*i*_, and each transition is the product of all the invasions happening between *τ*_*i*_ and *τ*_*i*+1_. For a fixed rate *ρ*, we modify the assembly graph *G*_*ρ*_ := *G*(*ρ*). Edges in *G*_*ρ*_ represent this new type of transition, and are weighted by the probability that such a transition occurs (e.g., 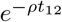 our example above). The vertices remain the feasible subsets. Define *G*_0_ = *G*_asym_; then, all edges in *G*_0_ appear for any *G*_*ρ*_ with non-zero probability. Increasing *ρ* will cause assembly to be able to access, with high enough probability to be observed, only feasible states with “fast” dynamics. Thus, at some value of *ρ*, we would only be able to access, if any, only the fixed point attractors for the top-down assembly. Fig. A1 shows how the fraction of feasible states observed along assembly changes as a function of *ρ*.

**Fig. A1:**
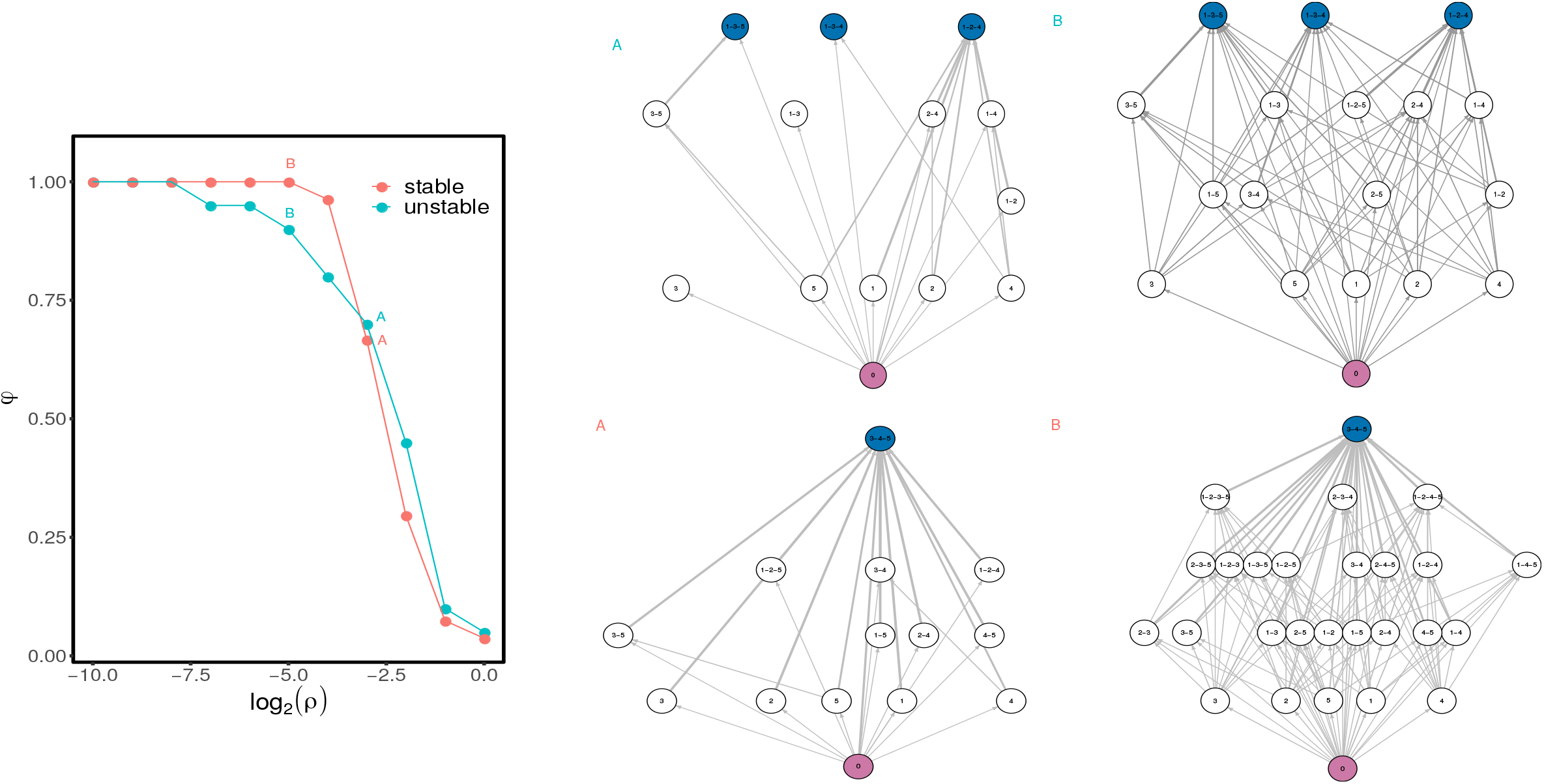
Fraction of accessible states as a function of the rate of invasion *ρ* for 500 random sequences of invasions of length 100, for the stable and unstable cases showed in Fig. 2 and 3 of the main text. On the right, we show the assembly graphs constructed under two particular rates. The width of the arrows represents the number of times this transition was observed during the simulations. Notice that the appearance of new sinks is due to the fact that we only consider as nodes states that reached their asymptotic equilibrium. Thus, the new sinks correspond to cases in which, although invasions are possible, a new equilibrium state is never reached due to the high invasion rate.

### S4 Symmetric Competitive Lotka-Volterra

In this section, we state the properties of the competitive GLV model studied in the main text, which allow us to conclude that the system satisfies all the assumptions of Theorem 1. Recall the equations:

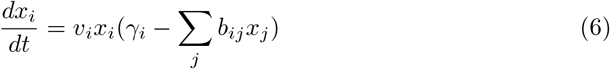

and assume that all parameters are positive. Additionally, assume that for *B* = (*b*_*ij*_), any principal submatrix *B*^*S*^ induced by a given set of species *S* is nonsingular, and that the matrix (*B*^*S*^ | *γ*^*S*^), for *γ*^*S*^ the subvector of growth rates induced by the species in *S*, has always full column rank.

MacArthur (1970) showed that, regardless of the stability properties of *B*:

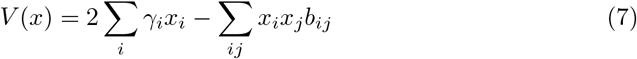

is maximized through the dynamics, and is thus a global Lyapunov function for the system. More explicitly, we find:

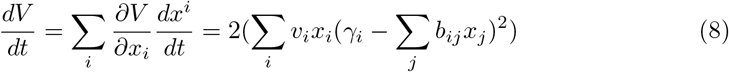

It then follows that either the system diverges, or the attractors are fixed points of the system—which are local maxima for *V* constrained on *x*_*i*_ *≥* 0. Because *b*_*ij*_ *>* 0, the first condition is impossible (Hofbauer & Sigmund, 1998), and thus all the attractors are constrained local maxima. We now show that the local maxima are given by fixed points *x*^*S*^, with positive entries only for the species present in *S*, and for which two additional properties hold:

a. *x*^*S*^ is non-invasible, i.e., for any *k ∉ S* we have:

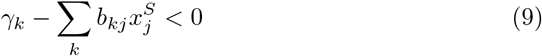
b. Let *B*^(*S*)^ be the submatrix of *B* containing the rows and columns only for the species present in *S*. Then *B*^(*S*)^ is positive definite.

Let *x*^*S*^ be a constrained local maximum for *V*, with 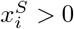 only for *i ∈ S*. Then, for any *k ∉ S* we must have:

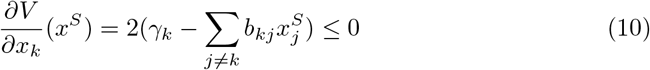

By the conditions on *B* and *γ*, the inequality is strict.

Now, let *V* ^*S*^ be *V* restricted to the species in *S*, and *y*^*S*^ be the restriction of *x*^*S*^ to *S*. Then *y*^*S*^ is an interior local maximum for *V* ^*S*^. It follows that the Hessian matrix of *V* ^*S*^, evaluated at *y*^*S*^, must be negative definite. A direct computation shows:

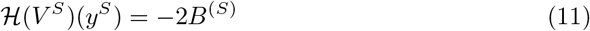

and thus *B*^(*S*)^ is positive definite.

Now consider a fixed point *x*^*S*^ as described above. Then, for any other vector *u* such that *x*^*S*^ + *u ≥* 0, i.e., *u*_*k*_ *≥* 0 for any *k ∉ S*, we have:

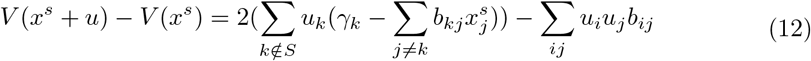

Because the second term is quadratic in *u*_*i*_, and the first sum is negative for any *u* with at least one *u*_*k*_ *>* 0, then taking |*u*_*i*_| *< ε* for some *ε* « 1 shows that *V* (*x*^*S*^ + *u*) *< V* (*x*^*S*^). For *u* with all *u*_*k*_ = 0, the result follows as above by the Hessian of *V* ^*S*^. Taking the intersection of both neighborhoods of *x*^*S*^, the claim follows.

### S5 Assembly of Consumer-Resource model

Lotka-Volterra systems with an interaction matrix as in Eq. (2) arise naturally from consumer-resource models. Let *C*_*i*_ and *R*_*k*_ be a set of consumers and resources whose dynamics are defined by the MacArthur’s model (MacArthur, 1970):

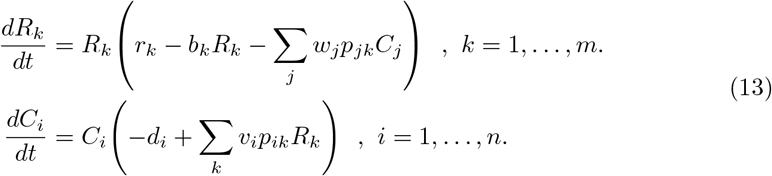

Where *v*_*i*_ can be understood as a consumer-specific efficiency of resource conversion, and *w*_*j*_ models the impact of consumers on resources. *d*_*i*_ and *r*_*k*_ are the mortality and growth rates of consumer and resources and *b*_*k*_ the intraspecific competition of resources. All parameters are positive. Assuming *R*_*k*_ is at equilibrium, then:

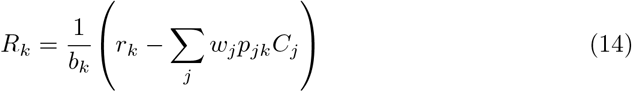

Thus replacing in the equation for the consumers we find:

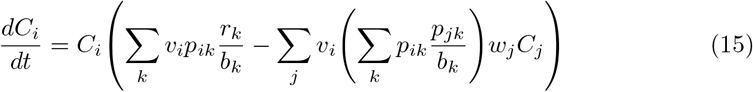

Letting 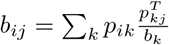, we recover Eq. (2) with *B* = *PD*(*b*)^−1^*P*^*T*^ positive definite for *m ≥ n*. The procedure implicitly assumes a separation of timescales between consumers and resources (with resources equilibrating faster than consumers), and the positivity of all resources at the given equilibrium. In what follows we show that the results of the main text will hold even if we analyze the full consumer-resource model under the assumption that the all the resources are always present (see also Case & Casten (1979); Marsland (2019); Mehta (2019)). Yet, relaxing the constraint on the coexistence of resources leads to different effects.

Let *d* = (*d*_*i*_), *r* = (*r*_*k*_), *γ* = (*r*, −*d*) and *P* = (*p*_*ik*_). Define *A* as:

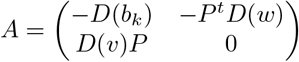

Let *N* = (*R, C*). Then, we recover the generalized Lotka-Volterra equations:

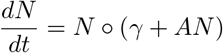

Where stands for the Hadamard (component-wise) product. By the special structure of *A*, it follows that it is a *B*-matrix (Hofbauer & Sigmund, 1998), thus all solutions to the system are uniformly bounded and there exists a saturated rest point. In (Case & Casten, 1979) it is stated that besides some degenerate cases the saturated rest point is the unique attractor of the dynamics. To avoid the degenerate cases it is enough to consider the following:

a. **Uniqueness of equilibrium solution** Any matrix submatrix of *P* has full column rank. Thus for any subsystem with number of resources at least as big as the number of consumers the equilibrium is unique.
b. **Strict invasibility** Let *Ã* be a principal submatrix of *A* induced by a subset of species *S*, and 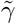 the subset of *γ* induced by the same set *S*. Then the matrix 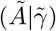 has full column rank. A consequence of this is that we cannot have an equilibrium with more predators than resources: If that were the case, then the induced *Ã* is singular and 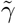 is in the span of *Ã* thus 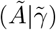 is not full column rank.

By performing a linear change of variables *C*_*i*_ *→ P*_*i*_ = (*w*_*i*_*/v*_*i*_)*C*_*i*_, *R*_*k*_ *→ Y*_*k*_ = *b*_*k*_*R*_*k*_, and *a*_*ij*_ = *v*_*i*_*p*_*ik*_*/b*_*k*_ we map the system to:

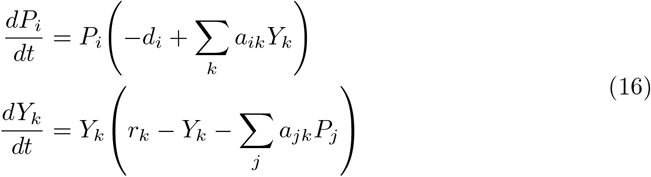

In this form, the results of (Mehta *et al*., 2019) say that the globally attractive fixed point minimizes the function:

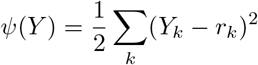

with the constraints:

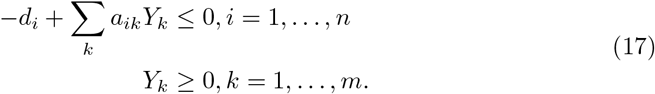

This follows by the application of the Karush-Kuhn-Tucker(KKT) conditions (Mehta *et al*., 2019): At the minimum points we have:

a. *P*_*i*_(−*d*_*i*_ + ∑*k a*_*ik*_*Y*_*k*_) = 0, for *i* = 1, *…, n*
b. *P*_*i*_ *≥* 0, for *i* = 1, *…, n*.
c. *u*_*k*_*Y*_*k*_ = 0, for *k* = 1, *…, m*.
d. *u*_*k*_ *≥* 0, for *k* = 1, *…, m*.
e. *r*_*k*_ − *Y*_*k*_ − *∑*_*j*_ *a*_*jk*_*P*_*j*_ = −*u*_*k*_ *≤* 0, for *k* = 1, *…, m*.

It then follows that any set of points satisfying the KKT conditions is a saturated rest point, thus by uniqueness the global attractor is the unique one satisfying them. By convexity of *ψ* the KKT conditions are sufficient, thus the globally attractive solution is the unique minimum.

The above framework can be used to study an assembly process in which the initial state is the state with all resources present and the consumers never cause extinction of the resources. In this way, we study only the assembly of the consumers in a changing background of density, but not identity, of resources. The claim is simply that *ψ* strictly increases along assembly: Indeed, let *S* = (*S*_*P*_) be a set of consumers. Choose a successful invader *j*, taking the system *S → S*^*′*^. By the non-invasibility conditions of the attractor it must be present in *S*^*′*^. Since *j* is a predator, the new equilibrium *Y* ^*S*^*′* minimizes *ψ* with an additional constraint, therefore *ψ*(*Y* ^*S*^*′*) *≥ ψ*(*Y* ^*S*^). By the uniqueness of the minimum and the invasibility condition we must have *ψ*(*Y* ^*S*^*′*) *> ψ*(*Y* ^*S*^) (see also discussion in (Marsland *et al*., 2019)). Thus by *ψ* we can order all the feasible equilibria of the system, thus by Lemma 1, the assembly graph would be acyclic. By the non-invasibility characterization of the attractors, Theorem 1 applies.

It is important to remark the importance of the assumption on the coexistence of resources. If the consumers can send resources extinct, we can find assembly processes with cycles, thus there does not exist an ordering of the states of the system. This comes from the fact that *ψ* actually decreases under successful invasions of resources: We can view *ψ* of the recipient community as a minimization problem with the additional constraint of *Y*_*k*_ = 0 for some *k*, while when we add the resource *Y*_*k*_ *≥* 0. By the invasibility condition of the new resource, the previous solution is no longer a minimum and thus *ψ* decreases. Fig. A2 shows assembly graphs for consumer resource systems in which we can find cycles, and we have that either it is still possible to reach the attractor of top-down assembly or not. In case a cycle exists, there is always the possibility that even if there is a way out of it the sequence of invasions is such that assembly does not take the system out of the cycle. Yet, if we assume that each successful invasion happens with the same probability, the probability of this scenario vanishes asymptotically. Thus, we expect that asymptotically we only observe cycles if they happen to be the endpoints of assembly (right panel in fig Fig. A2).

**Fig. A2:**
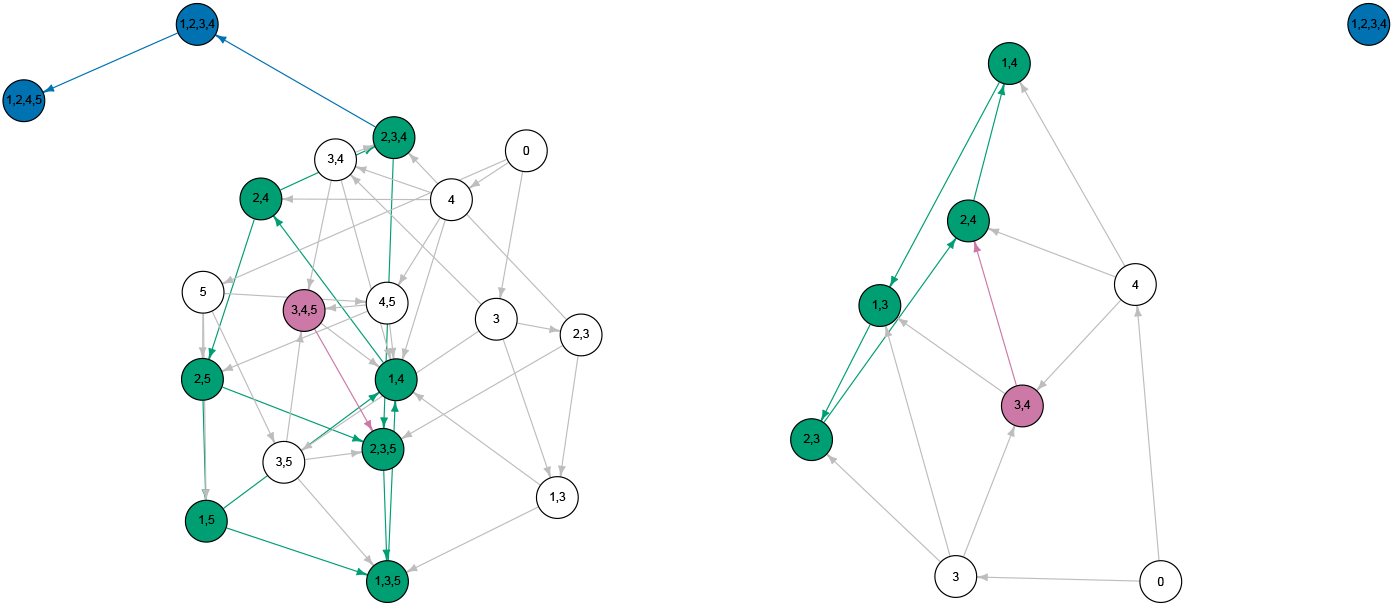
Assembly graph for consumer resource systems in which predators can make resources go extinct. In this case we see that cycles are not precluded: A strongly connected component for each graph is highlighted in green. Yet, we can have *G* accessible (left) or not (right). A path from the cycle to the top-down assembly attractor, if it exists, is highlighted in blue. The state with all resources present and an edge taking it to the respective cycle is highlighted in violet.

### S6 Construction of G

The construction of the assembly graph, even in the simple setting of a GLV model, is a demanding computational task. For a set of parameters (*A, r*), in order to define the nodes of *G* we need to compute the set of all feasible sub-communities. The number of possible sub-communities is bounded by 2^*n*^ from above, and its actual size depends on the parameters. Nevertheless, if the size of feasible sets is a non-vanishing fraction of the total number of possible communities, we have exponential growth of the size of the graph with *n*. This makes the construction of the assembly graph for large enough communities computationally intractable. Regardless of this constraint, we believe that *G* could be helpful to visualize assembly for small (*<* 30 species) communities under experimental conditions, or as an arena where new theoretical devices can be applied to better understand assembly.

